# Maternal dietary deficiencies in folic acid or choline reduce primary neuron viability after exposure to hypoxia through increased levels of apoptosis

**DOI:** 10.1101/2023.12.09.570894

**Authors:** Alice Yaldiko, Sarah Coonrod, Purvaja Marella, Lauren Hurley, Nafisa M. Jadavji

## Abstract

Stoke is the leading cause of death and disability globally. By addressing modifiable risk factors, particularly nutrition, the prevalence of stroke and its dire consequences can be mitigated. One-carbon (1C) metabolism is a critical biosynthetic process that is involved in neural tube closure, neuronal plasticity, and cellular proliferation in the developing embryo. Folic acid and choline are two active components of 1C metabolism, we have previously demonstrated that maternal dietary deficiencies in folic acid or choline worsen stroke outcomes in offspring. However, there is insufficient data to understand the neuronal mechanisms involved. We exposed embryonic neurons of offspring, whom mothers were on folic acid or choline deficient diets, to hypoxia conditions for 6 hours and return to normoxic conditions for 24 hours to model an ischemic stroke and reperfusion injury. To determine whether increased levels of either folic acid or choline can rescue reduced neuronal viability, we supplemented cell media with folic acid and choline prior to and after exposure to hypoxia. Our results suggest that maternal dietary deficiencies in either folic acid or choline during pregnancy negatively impacts offspring neuronal viability after hypoxia. Furthermore, increasing levels of folic acid or choline prior to and after hypoxia have a beneficial impact on neuronal viability. The findings contribute to our understanding of the intricate interplay between maternal dietary factors, 1C metabolism, and the outcome of offspring to hypoxic events, emphasizing the potential for nutritional interventions in mitigating adverse outcomes.

## Introduction

Stroke is the leading cause of death and disability worldwide [1], [2]. It is commonly known for its detrimental effects such as paralysis of the limbs and reduced ability to communicate [2]. Stroke presents in multiple forms; seventy percent of strokes have been ischemic. Ischemic stroke occurs when there is a sudden loss of blood flow to a particular area of the brain, leading to tissue loss and neurological problems which have the potential to be irreversible [3]. The population most susceptible to stroke is individuals over the age of 65; however, stroke can present at any age [4]. The proportion of world’s population above the age of 65 years has increased in the last few years correlating with a rise in the prevalence of stroke, creating a heightened need to address the issue [5]. Not only age but other risk factors, such as increased triglycerides, cholesterol, and diabetes, contribute to increased risk of stroke [6]–[8]. Fortunately, due to the influence of diet on some of the aforementioned risk factors, the incidence of developing stroke can be decreased by modifying nutrition [9].

An example of nutritional link to ischemic stroke is one-carbon (1C) metabolism which integrates nutritional signals with several biosynthetic processes, redox homeostasis, and epigenetics [10]–[12]. The nutrient, choline and B-vitamin, folic acid, are central to 1C. Genetic deficiencies in 1C has shown a direct impact on neuronal and astrocyte viability, and thus worsened stroke outcomes [13], [14]. Although the mechanism for this relation is still unclear, one common association is that folic and choline deficiency results in hyperhomocysteinemia, a common risk factor for developing ischemic stroke [15]. Both, folic acid and choline are important for the closure of the neural tube *in utero* [16]. Choline also plays an important role in lipid metabolism and acetylcholine synthesis which are essential during neurodevelopment [17]. Thus, deficiencies in these two components can lead to defective neurological development and consequently increased risk of stroke. On a positive note, 1C metabolites supplementation after stroke may be responsible for plasticity and brain recovery [13], [18]–[20]. As food fortification programs and the encouragement of supplemental prenatal vitamins becomes more prevalent, it is important to develop a better understanding of the relation between maternal nutrition and offspring health, particularly stroke outcome. Maternal nutrition during pregnancy and lactation is recognized as a critical factor determining the health of offspring. The developmental origins of health and disease (DOHaD) theory suggests that prospective chronic diseases are programmed *in utero*-giving rise to programming of offspring cardiovascular, metabolic, and neuroendocrine dysfunction [21]– [23]. The aim of this study was to investigate the impact of maternal folic acid or choline deficiencies on offspring primary neuronal viability after hypoxia, an *in vitro* model of stroke, as well as the role of supplementation of choline or folic acid.

## Materials and Methods

All experiments were conducted in accordance with animal welfare regulations, and the protocols were approved by the Midwestern University IACUC Committee (Protocols 2977 and 2978). Experiments were performed in a fully randomized and blinded fashion.

### Primary Neuron Isolation

Female and male wildtype mice were housed together for 24 h after which embryos were collected on day 17 [13]. Neurons were isolated from cortical tissue using the papain dissociation kit (Worthington Biochemical Inc.) as previously described [24] [25] and cells were plated at a density of 175,000 cells/cm^2^. Cultures were prepared with supplemented neurobasal media (Life Technologies) in 24-well plates as previously described [26]. Cells were grown in culture for 8 days prior to hypoxia treatment [13], [27].

### Hypoxia: In vitro model of stroke

Neurons remained in medium and were incubated in ∼1% O_2_ (5% CO_2_, 37°C) for 360 min in a triple-gas incubator (Eppendorf) and then returned to normoxic conditions (21% O_2_, 5% CO_2_, 37°C) [28]. Control plates remained in normoxic conditions for the same amount of time. Twenty-four hours after hypoxia damage cells were assessed for cell viability or fixed with 4% PFA and used for immunofluorescence staining. All cell culture experiments, including cell viability and immunofluorescence studies were repeated 5 times using different pregnant females for each experiment [13], [27].

### Impact of maternal folic acid or choline dietary deficiency on offspring primary neurons after hypoxia

Experimental manipulations are summarized in Figure 1A. Two-month-old C57Bl/6 female mice were obtained from Jackson Laboratory and habituated for seven days before they were placed on either control (CD, folic acid 3 mg/kg and choline bitartrate, 1150mg/kg), folic acid (FADD, folic acid, 0.3 mg/kg), or choline (ChDD, choline bitrate 300 mg/kg) deficient diets. The dams were maintained on the diets 4 weeks prior to and during pregnancy. Female and male wildtype mice were housed together for 24 h after which embryos were collected on day 17 for primary neuronal cultures and hypoxia treatment [13].

**Figure 1.**
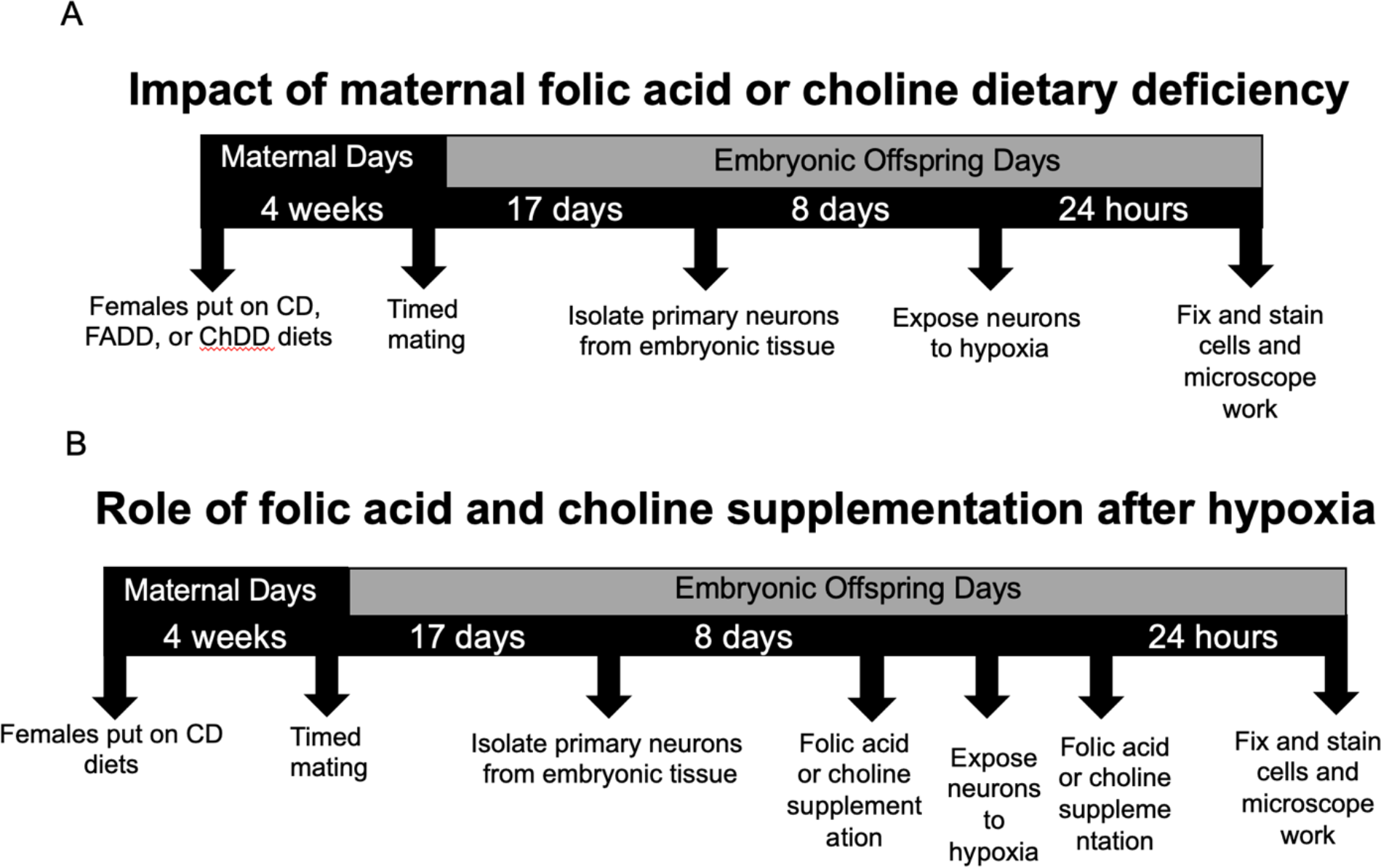
Experimental timeline for *in vitro* stroke experiments. For impact of maternal folic acid or choline dietary deficiency study female mice were placed on one of 3 diets: folic acid deficient (FADD), choline deficient (ChDD), or control diet (CD). This is done for 4 weeks prior to mating. The females will be mated with the males for a period of 24 hours. At embryonic day 17 primary cortical neurons will be isolated from embryos. These neurons are grown in culture for a period of 7 days. On day 8, the cells are exposed to hypoxic conditions (∼1% O_2_) for 6 hours using a hypoxia chamber and then returned to normal O_2_ condition for 24hr (∼21% O_2_). To determine the role of folic acid and choline supplementation after hypoxia on neuronal viability post stroke, folic acid (250mg/ml) or choline chloride (250mg/ml) were added to primary neuronal cells prior to and after exposure to hypoxia. Primary neurons are fixed using paraformaldehyde and stained with trypan blue staining to assess cell viability. Microscopy was used to visualize saining and NIH ImageJ was used to quantify living and dead cells.

### Impact of folic acid and choline supplementation on primary neurons after hypoxia

Experimental manipulations are summarized in Figure 1B. Neurons from CD fed female embryos were incubated with 250 mg/ml of folic acid or choline chloride immediately before and after exposure to hypoxia [29], [30].

### Cell viability

Twenty-four hours after hypoxia damage, neuronal viability was assessed using trypan blue staining [13], [27], [31]. Cultures were imaged using Olympus IX73 inverted florescent microscope, images were taken at 200X. The number of alive and dead cells were counted by two individuals blinded to experimental groups.

### Immunofluorescence

We investigated apoptosis in primary neurons from maternal deficiency experiment. Neurons were incubated with anti-active caspase-3 (1:100, Cell Signaling Technologies, RRID: AB_331439) which was diluted in 0.5% Triton X overnight at 4°C. The following day, neurons were incubated in secondary anti-Rabbit Alexa Fluro 555 (Cell Signaling Technologies, RRID: AB_10696896) at room temperature for 2 h and then stained with 4’, 6-diamidino-2-phenylindole (DAPI) (1:10000, Thermo Fisher Scientific). Semi-quantitative methods were utilized for analysis, co-localization of active caspase-3 with DAPI were counted and averaged per well. Staining was visualized using the Olympus IX73 inverted florescent microscope and all images were collected at the magnification of 40X. All cell counts were performed by 2 individuals blinded to the experimental treatments.

## Statistical analysis

Microscopy analysis was conducted by two individuals blinded to experimental groups. Data from the study were analyzed using GraphPad Prism 10.0. Two-way ANOVA analysis was performed when comparing the mean measurement of treatment group for the cell viability and *in vitro* immunofluorescence staining. Main effects of two-way ANOVAs were followed up with Tukey’s post-hoc test to adjust for multiple comparisons. All data are presented as mean + standard error of the mean (SEM). Statistical tests were performed using a significance level of 0.05.

## Results

### Maternal dietary deficiencies in folic acid or choline reduce cell viability in primary neurons from offspring exposed to hypoxia

To confirm maternal diet did not impact neuronal viability under normoxic conditions, we measured living and dead cells using Trypan blue. Representative images from all groups are shown in Figure 2A. There were no differences between maternal dietary groups in the percent living (Figure 2B, F (_2, 15_) = 0.189, p = 0.83) and dead (Figure 2C, F (_2, 13_) = 0.710, p = 0.51) cells.

**Figure 2.**
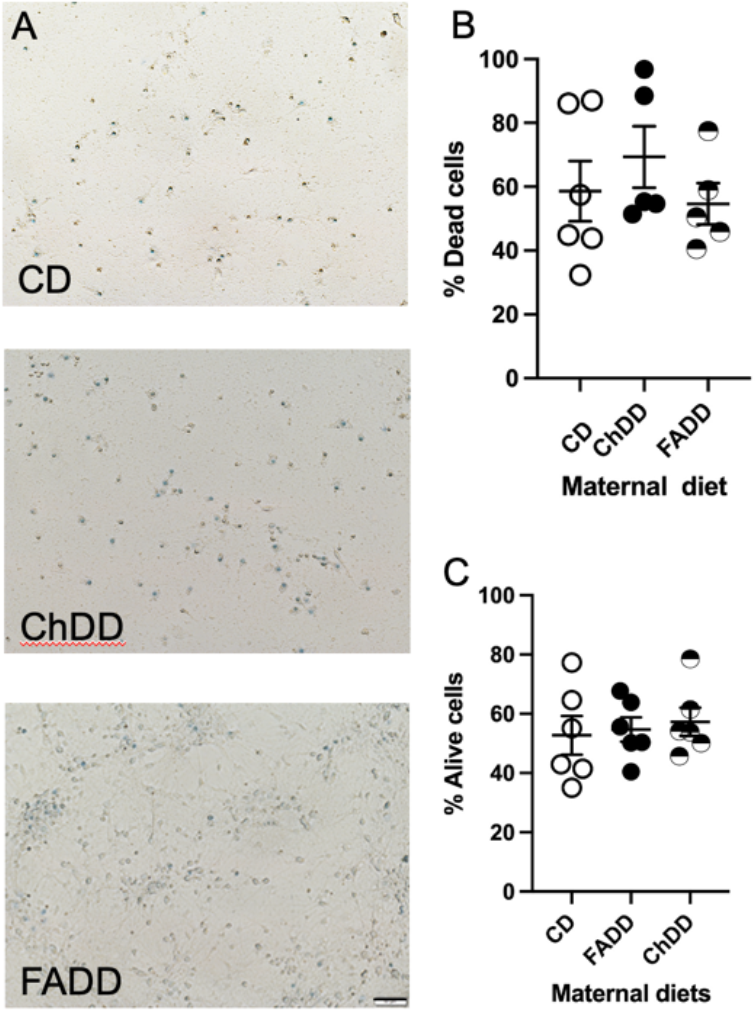
Cell viability of primary neurons of offspring maintained on folic acid (FADD), choline deficient (ChDD), and control (CD) diets under normoxic conditions. A) Representative images of trypan blue staining for control plate. The percentage of dead cells B) and alive cells C) per each maternal diet when exposed to normoxic conditions. There are 5 to 6 females per group. Scale bar at 50μm.

When we exposed primary neuronal cells to hypoxia, we observed a difference between maternal dietary groups in the cell viability, representative images are shown in Figure 3A. There were reduced living cells because of maternal diet (Figure 3B, F (_2, 13_) = 6.61, p = 0.01). There were reduced living neurons from embryos from ChDD mothers (p = 0.05) when compared to control embryos. Furthermore, there were more dead neurons from maternal deficient diet in either folic acid or choline (Figure 3C, F (_2, 15_) = 8.12, p = 0.0041). There were increased dead neurons from FADD (p = 0.01) and ChDD (p = 0.05) mothers when compared to controls.

**Figure 3.**
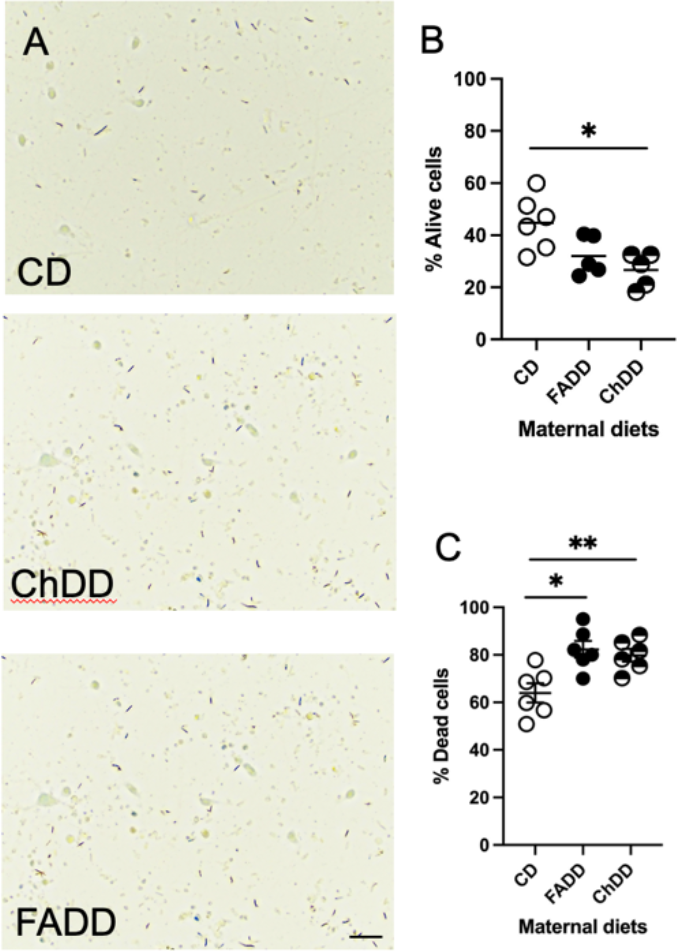
Cell viability of hypoxic primary neurons of offspring maintained on folic acid (FADD), choline deficient (ChDD), and control (CD) diets. A) Representative images of trypan blue staining for hypoxic plate. The percentage of dead cells B) and alive cells C) per each maternal diet when exposed to hypoxia. There are 5 to 6 females per group. Scale bar at 50μm. * p < 0.05, ** p < 0.01 Tukey’s pairwise comparison.

### Both hypoxia and maternal diet increase the number of apoptotic cells

We measured apoptosis by employing a semi-quantitative method. We counted the number of active caspase-3 co-localized with DAPI from the three different maternal diets after hypoxia treatments. Representative images are shown in Figure 4A. Both hypoxia (Figure 4B, F(_1,24_) = 892.4, p < 0.0001) and maternal diet (F (_2,24_) = 36.65, p < 0.0001) increased the number of active caspase-3 positive cells.

**Figure 4.**
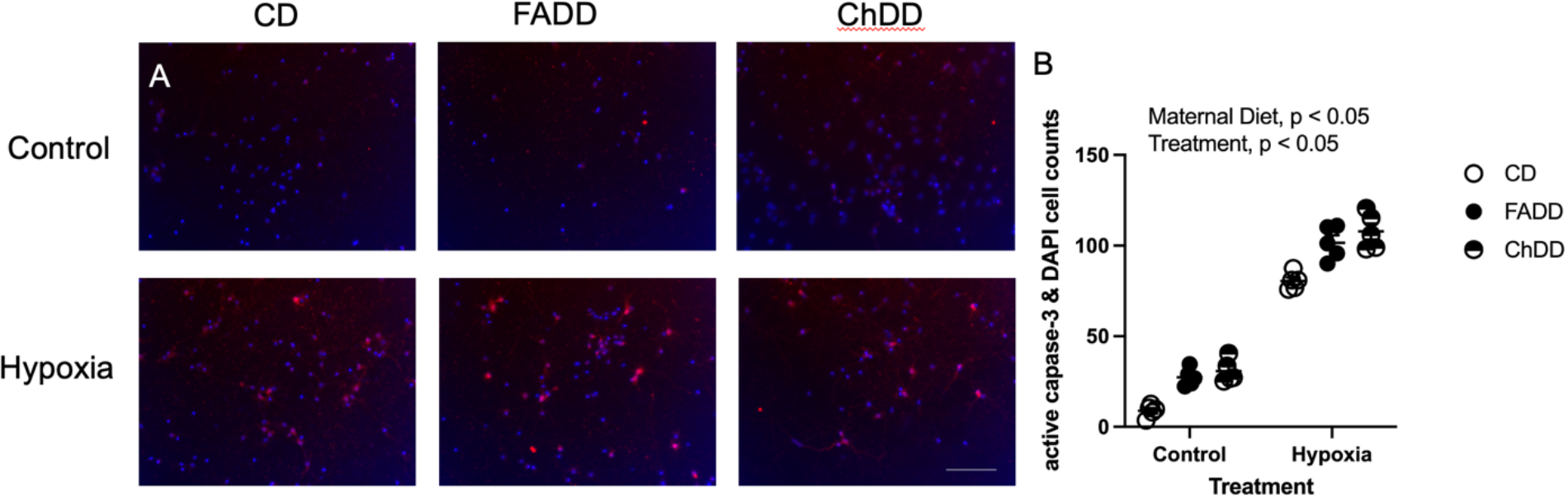
Apoptosis in primary neuronal culture of offspring from mothers maintained on folic acid (FADD), choline deficient (ChDD), and control (CD) diets using an immunofluorescent staining. Representative images of the (A) CD (B) FADD and (C) ChDD for the control plate and hypoxia plate. D) Quantification of apoptotic neurons per each maternal diet when exposed to normoxic conditions and hypoxic conditions. There are 4 to 5 females per group. Scale bar at 50 μm.

### Impact of folic acid and choline supplementation on primary neurons after hypoxia

Since a maternal deficiency in either folic acid or choline reduced cell viability and increased apoptosis in offspring primary neurons, we assessed the impact of supplementation of folic acid or choline on neuronal viability before and after hypoxia. Representative images of Trypan blue staining can be seen in Figure 5A. After hypoxia, there was no impact of supplementation on the dead cells (Figure 5B, F (_2,23_) = 0.92, p = 0.41), but there were more living cells in supplemented folic acid or choline media (Figure 5C, F (_2,14_) = 15.20, p = 0.0003). Both folic acid (p = 0.0027) and choline (p = 0.0004) increased the number of alive cells compared to nontreated cells.

**Figure 5.**
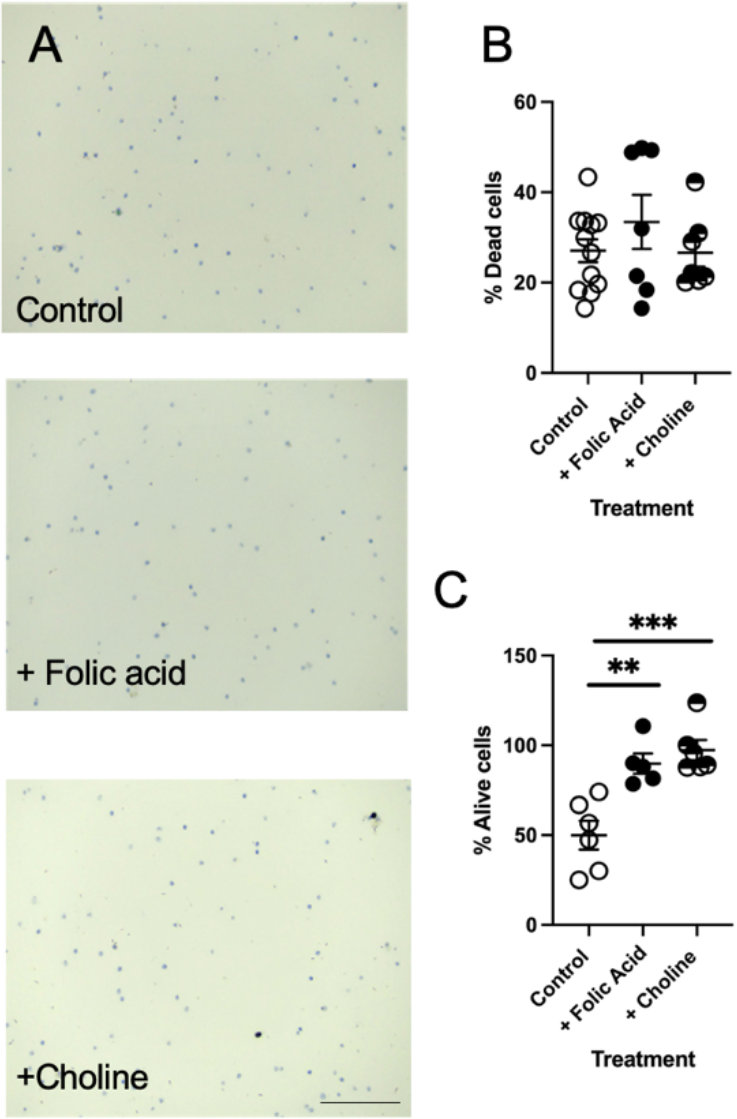
Cell viability of primary neurons with folic acid and choline chloride supplementation prior to after hypoxia. A). Representative images of trypan blue staining for control plate. Quantification of the percentage of dead B) and living C) cells per each maternal diet when exposed to normoxic conditions. There are 5 to 10 females per group. Scale bar at 50μm. ** p < 0.01, *** p < 0.001, Tukey’s pairwise comparison.

## Discussion

Stroke is the leading cause of death and disability globally [2]. Through modifiable risk factors, such as nutrition, stroke’s prevalence and the devastating outcomes can be minimized [32], [33]. One-carbon (1C) metabolism is a critical biosynthetic process that is involved in the closure of the neural tube, neuronal plasticity, and cellular proliferation in the developing embryo. We have previously demonstrated that maternal dietary deficiencies in folic acid or choline worsen stroke outcomes in offspring[34], [35]. However, there is insufficient data to understand the mechanisms involved. For this study we exposed primary neurons of offspring from mothers deficient in either folic acid or choline to hypoxia. Offspring primary neurons from mothers deficient in folic acid and choline had reduced neuronal viability and increased levels of apoptotic cells after hypoxia. Interestingly, supplementation with folic acid and choline, both before and after exposure to hypoxia conditions, yielded an increase in the percent of alive neurons. These results indicate mechanisms through which 1C impacts neurons after hypoxia.

Offspring neuronal development is tightly linked to maternal nutrition [36]. It has been established that folic acid and choline are two important dietary factors that are involved in neuronal development [37]. There are numerous proposed mechanistic pathways in which these two molecules are believed to partake in; therefore, it is still unclear exactly how they function to decrease neural developmental defects [38]. Our results show that neurons from offspring whose mothers’ diets were deficient in folic acid or choline were challenged under hypoxic conditions. Folic acid is an essential molecule when it comes to nucleotide synthesis. Nucleotides are the building blocks of DNA. For cellular proliferation to take place, cells need to synthesis DNA. Folic acid is used in *de novo* purine biosynthesis and *de novo* deoxythymidine monophosphate synthesis [38]. Methylation is another pathway in which these molecules contribute. Folic acid and choline are involved in homocysteine breakdown pathway via remethylation. *S*-adenosylmethionine (SAM) is known as the universal methyl donor, it serves as the substrate in many methylation reactions [38]. Upon donating a methyl group, SAM is converted to S-adenosylhomocysteine (SAH); subsequently, SAH is converted to adenosine and homocysteine [38]. Here folic acid and/or choline’s role comes to play in methylating homocysteine. If there is a deficiency in folic acid or choline, accumulation of homocysteine is observed and the reaction favors the formation of SAH which is known to be a potent methyltransferase inhibitor [38]. The accumulation of homocysteine may lead to the activation of p53 which is a tumor suppressor gene [39]. This may result in increased apoptosis and can possibly explain the increased levels of apoptosis we observed. Through 1C, folic acid and choline are involved in numerous metabolic pathways; thus, it is safe to suggest that sufficient amounts are needed for cellular neurodevelopment.

Supplementation with folic acid and choline before and after hypoxia increased neuronal viability. However, it is important to note that there was no difference in the percent of dead cells. There are numerous proposals explaining how supplementation with folic acid or choline increases cellular viability. A simple explanation is supplementing with these two molecules can help overcome the increased requirement during cellular stress by merely increasing their bioavailability [38]. A study also found that women with neuronal tube defect fetuses had antibodies targeted against folic acid receptors blocking its uptake [38]. A separate study showed that increasing folic acid supplementation after cellular injury resulted in an increased number in folic acid receptors [39]. Furthermore, as explained previously, folic acid and choline are involved in homocysteine breakdown pathway. Increased levels of homocysteine can result in oxidative stress which increases the risk of stroke [39]. We have previously shown an increase in neuronal plasticity and reduction of homocysteine after exposure to ischemic stroke *in vivo* when mice were supplemented with folic acid and choline, as well as other 1C metabolites [39]. Our previous study also showed a decreased number of apoptotic cells which can be linked to the relationship between p53 and homocysteine, as explained earlier [39].

Maternal nutrition during pregnancy and lactation is recognized as a critical factor determining the health of offspring [1]. The DOHaD theory suggests that prospective chronic diseases are programmed *in utero*-giving rise to programming of offspring cardiovascular, metabolic, and neuroendocrine dysfunction [21], [22]. This study has shed light on the impact of maternal dietary folate and choline deficiencies on offspring primary neuronal function after hypoxia. We think the next steps would be to assess the impact of maternal dietary deficiencies on glial cell function [27] and whether supplementation with folic acid or choline supplementation are more beneficial. Since 1C metabolism plays an important role in the generation of SAM, we think it would be worthwhile to investigate methylation level and gene expression of key genes involved in 1C and hypoxia outcome in offspring.

## Acknowledgments

None

## Data Availability Statement

The data that supports the findings of this study are available in the supplementary material of this article.

## Notes on Contributors

Alice Yaldiko, BS is a Doctor of Osteopathic Medicine student at Midwestern University.

Purvaja Marella, BS is a Doctor of Osteopathic Medicine student at Midwestern University.

Sarah Coonrod, BSc is a Doctor of Veterinary Medicine student at Midwestern University.

Nafisa M. Jadavji PhD is an Assistant Professor in Biomedical Sciences at Midwestern University (US) and Research Assistant Professor in Neuroscience at Carleton University (Canada).

